# Intermittent fasting induces rapid hepatocyte proliferation to restore the hepatostat in the mouse liver

**DOI:** 10.1101/2021.10.16.464650

**Authors:** Abby Sarkar, Yinhua Jin, Brian C. DeFelice, Catriona Logan, Yan Yang, Teni Anbarchian, Peng Wu, Maurizio Morri, Norma Neff, Huy Nguyen, Eric Rulifson, Matt Fish, Azalia M. Martínez Jaimes, Roel Nusse

## Abstract

Nutrient availability fluctuates in most natural populations, forcing organisms to undergo periods of fasting and re-feeding. It is unknown how dietary changes influence liver homeostasis. Here, we show that a switch from ad libitum feeding to intermittent fasting, IF, promotes rapid hepatocyte proliferation. Mechanistically, IF-induced hepatocyte proliferation is driven by the combined action of intestinally produced, systemic endocrine FGF15 and localized WNT signaling. Hepatocyte proliferation during periods of fasting and re-feeding re-establishes a constant liver-to-body-mass ratio, thus maintaining the hepatostat. This study provides the first example of dietary influence on adult hepatocyte proliferation and challenges the widely held view that liver tissue is mostly quiescent unless chemically or mechanically injured.

## Introduction

Periods of fasting and re-feeding induce profound tissue remodeling and regeneration in several tissues including the intestine (O’Brien, Soliman, Li, & Bilder, 2011; Yilmaz et al., 2012), the muscle (Cerletti, Jang, Finley, Haigis, & Wagers, 2012) and blood (Brandhorst et al., 2015; J. Chen, Astle, & Harrison, 2003; Ertl, Chen, Astle, Duffy, & Harrison, 2008). These tissue changes are thought to be mediated through diet-induced growth factor signaling, including both local (paracrine) and systemic (endocrine) signals that influence cell biology and function (Mihaylova, Sabatini, & Yilmaz, 2014). The impact of fasting and re-feeding on liver tissue homeostasis is unknown.

In contrast to other organs, the liver maintains a constant ratio with body weight to preserve homeostasis—this is termed the hepatostat (Michalopoulos, 2021). For example, when injured, the liver restores this ratio through hepatocyte renewal, resulting in the liver’s ability to maintain its many metabolic functions that are executed by hepatocytes. Organized into hexagonal lobular units, hepatocytes are stacked in between a central vein and a portal triad, that consists of a portal vein, hepatic artery, and bile duct. The directional flow of oxygenated blood from the portal to central axis creates a gradient of cytokines, nutrients and growth factors throughout the liver lobule that influences hepatocyte transcriptome and function (Halpern et al., 2017). Thus, pericentral hepatocytes, present near the central vein, receive different growth factor signals compared to the mid-lobular and periportal hepatocytes that occupy the rest of the liver lobule. Hepatocyte turnover along the liver lobule has been well characterized during ad libitum (AL) feeding (F. Chen et al., 2020; He et al., 2021; Lin et al., 2018; Wang, Zhao, Fish, Logan, & Nusse, 2015; Wei et al., 2021), when animals are given constant access to food. In these studies hepatocytes have an detectable rate of turnover and division (He et al., 2021; Wei et al., 2021), but the proliferation rates compared to other tissues are low (Michalopoulos, 2021). In contrast, no study so far has looked at hepatocyte turnover during periods of fasting and re-feeding, arguably a dietary state that more closely mimics nutrient availability and intake in natural populations, where periods of food availability fluctuate.

### Rapid proliferation of pericentral hepatocytes occurs during intermittent fasting

To determine the impact of IF on hepatocyte turnover, we compared the spatial expression of proliferation marker Ki67, in 1-week and 3-week IF-treated livers compared to AL-treated livers (Figure 1 A-C). We assessed the presence of Ki67+ hepatocytes throughout the liver lobule using a pericentral hepatocyte marker (Glutamine synthetase) and a periportal hepatocyte marker (E-cadherin) (Figure 1A). Total Ki67+ hepatocytes increased by 2-fold at 1-week of IF-treatment compared to AL-treatment (Figure 1B). This increase was no longer observed at 3-weeks of IF-treatment, suggesting that the increase in proliferation was short term (Figure 1B). The number of pericentral Ki67+ hepatocytes increased by approximately 120% after 1 week of IF treatment compared to AL treatment and 3-weeks of IF treatment (Figure 1C). At 3 weeks of IF treatment, the number of Ki67+ hepatocytes returned to AL levels, and were predominately midlobular (Figure 1C), as has been previously described (He et al., 2021; Wei et al., 2021).

**Figure 1.**
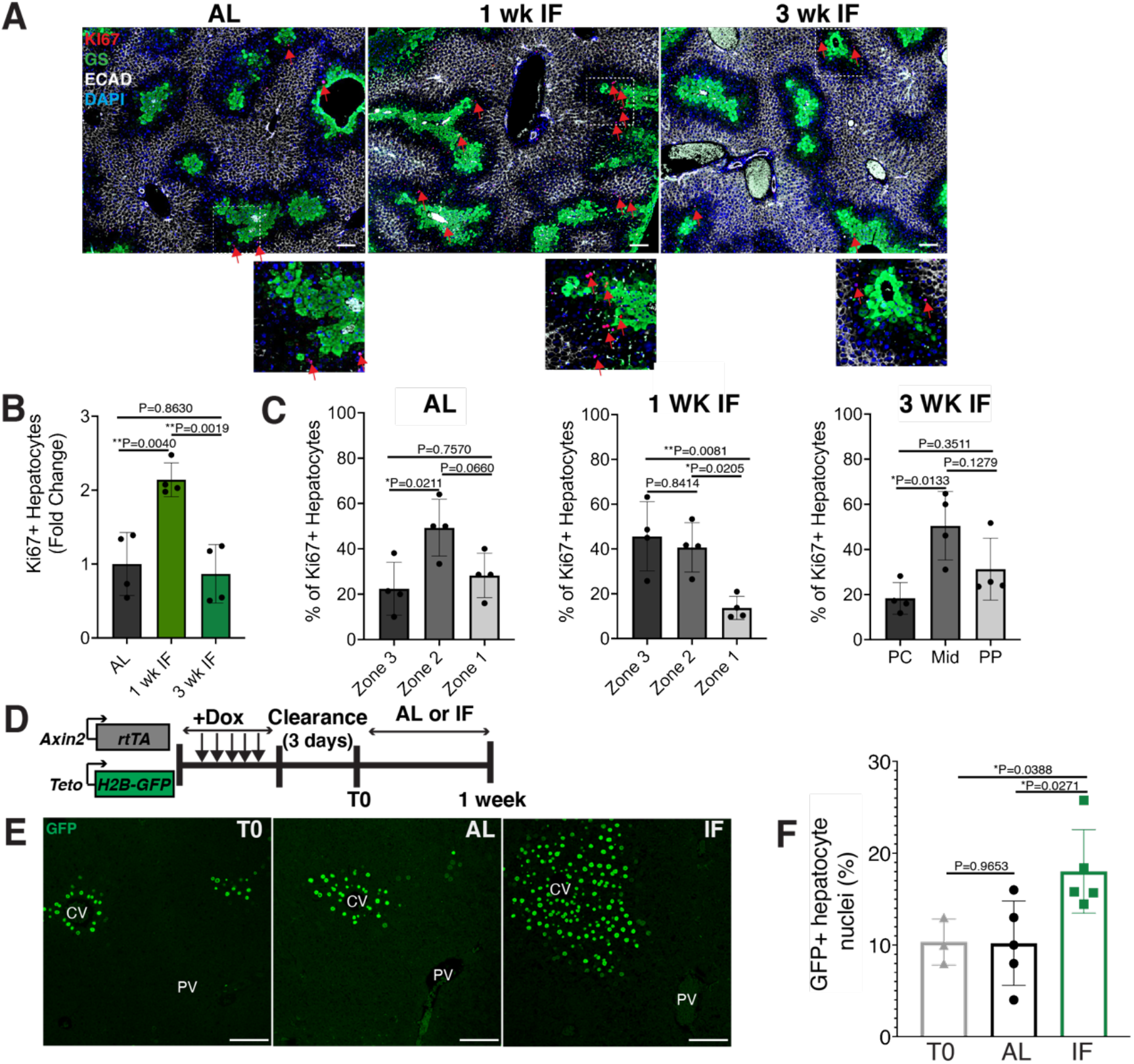
Intermittent fasting induces rapid hepatocyte proliferation. **(A)** Ki67 immunoflourescence for the detection of proliferating cells in AL, 1-week IF and 3-weeks IF treated livers. IF-treated livers were analyzed 30 minutes after re-feeding cycle. **(B-C)** Quantification of spatial distribution and percentage of Ki67+ hepatocytes in AL and IF treated livers. One-way ANOVA, N=4. **(D)** Doxycycline inducible *Axin2-rtTA*; *Teto-H2BGFP* system to label all pericentral hepatocytes and trace cell proliferation. Mice were pulsed with doxycycline (Dox) for 7 days, cleared of dox for 3 days and ad libitum fed or intermittently fasted for 6 days. **(E)** GFP immunofluorescent images showing increased hepatocyte expansion in AL and IF compared to TO. **(F)** Percentage of GFP+ hepatocyte nuclei in AL, IF livers from A. One-way ANOVA. N=3 (T0), 5 (AL), 5 (IF). \*\**P*<0.01; \**P*<0.05. Error bars indicate standard deviation. Scale bar, 100μm. wk, weeks.

Next, we corroborated the rapid and positional shift in IF-induced hepatocyte proliferation by employing both random (Supplementary Figure 1) and pericentral-specific cell lineage tracing systems (Figure 1 and Supplementary Figure 1) to capture representative hepatocytes and trace their clonal expansion in the liver under an AL or IF feeding regimen. First, to study hepatocyte proliferation throughout the liver lobule, we employed an inducible, *R26-CreERT2 (Ventura et al., 2007)* allele to permanently and randomly label cells with one of the four fluorophores in the *R26-Confetti* allele (Snippert et al., 2010). Following two weeks of tamoxifen clearance from the liver (time zero [T0], Supplementary Figure 1A), greater than 95% of labeled cells co-expressed one of the fluorophores and the hepatocyte specific transcription factor HNF4A (Supplementary Figure 1B), demonstrating labeling of mostly single hepatocyte clones distributed throughout the liver lobule (Supplementary Figure 1C). 1-3 weeks after IF feeding, a distinct increase in pericentral clone size was observed compared to AL fed animals (Supplementary Figure 1C, D). Second, to study pericentral hepatocyte proliferation kinetics during ad libitum feeding and intermittent fasting, we utilized an inducible system to mark and trace pericentral hepatocytes. In *Axin2-rtTA*; *TetO-H2B-GFP* transgenic mice (Tumbar et al., 2004; Yu, Liu, Costantini, & Hsu, 2007), a modified promoter of the WNT transcriptional target gene, *Axin2*, is used to control expression of a stable histone 2B-GFP fusion protein with doxycycline (dox) administration, thus marking WNT-responsive, pericentral hepatocytes and their progeny (8). Importantly, in these mice the ectopic *Axin2-rtTA* expression cassette leaves the endogenous *Axin2* locus unchanged. *Axin2rtTA; TetO-H2BGFP* animals were given doxycycline (dox) for 7 days, then cleared of dox for 3 days, and analyzed after dox clearance (T0) or after an additional 6 days of the AL or IF feeding regimen (Figure 1D). We observed a 74% increase in GFP-labeled hepatocyte nuclei in IF-treated animals compared T0 animals, thus confirming expansion of pericentral hepatocytes (Figure 1E, F). No significant change was observed between AL and T0 animals. Additionally, we quantified pericentral hepatocyte proliferation kinetics between 1 week, 3 weeks and 3 months of IF and AL-treatment (Supplementary Figure 1E-F). Notably, the majority of labelled hepatocyte expansion occurred within the first week of IF-treatment. Together these data demonstrate the rapid and transient proliferation of pericentral hepatocytes during IF feeding.

As an independent means to characterize hepatocytes after IF treatment, we performed single cell-RNA sequencing on livers from 1 week IF-treated and AL-treated animals. Livers were collected during a matched, neutral feeding and circadian state. Sequenced hepatocytes were classified into pericentral, mid-lobular and periportal hepatocytes (Supplementary Figure 2 A-B), using well characterized marker genes (Halpern et al., 2017).The proportion of hepatocytes enriched for pericentral transcripts in IF-treated animals was 19.52 ± 4.9%, more than twice as much as the proportion observed in AL-treated animals, 8.97 ± 1.2%. The increased proportion of pericentral hepatocytes in IF-treated animals suggests that IF increases the number of pericentral hepatocytes rather than mid-lobular or periportal hepatocytes. Furthermore, within pericentral hepatocytes, gene expression analyses revealed a distinct increase in *de novo* lipogenesis genes (*Fasn*, *Scd1* and *Acyl*) in IF compared to AL livers (Supplementary Figure 2 C), highlighting a cellular mechanism for the induction of de novo lipogenesis in the liver, a phenomenon previously observed during intermittent fasting (Hatchwell et al., 2020).

### Nutrient-responsive endocrine FGF15-β-KLOTHO signaling induces hepatocyte proliferation during intermittent fasting

Endocrine FGF signaling is critical in mediating an organism’s physiological response to fasting and re-feeding (Potthoff, Kliewer, & Mangelsdorf, 2012). Upon re-feeding, FGF15, produced by intestinal enterocytes, travels through the bloodstream and binds to its co-receptor, β-KLOTHO (KLB), on hepatocytes (Inagaki et al., 2005). Endocrine FGF signaling has also been shown to play important roles in regulating hepatocyte metabolism (Inagaki et al., 2005; Potthoff et al., 2011) and regeneration (Uriarte et al., 2013). To investigate this further, we conducted a kinetic and a histology screen to identify when and where FGF15 acts on hepatocytes during IF. In IF-treated animals, *Fgf15* expression was rapidly induced upon re-feeding, reaching maximal levels in the intestine within 30 minutes upon re-feeding (Figure 2A), compared to AL animals which did not detectably express *Fgf15* (Figure 2B). 30 mins after re-feeding, downstream pathway components KLB, PHOSPHO-TYROSINE and PHOSPHO-C-JUN were elevated in IF animals, and all of these three components were preferentially concentrated in the pericentral region (Figure 2B).

**Figure 2.**
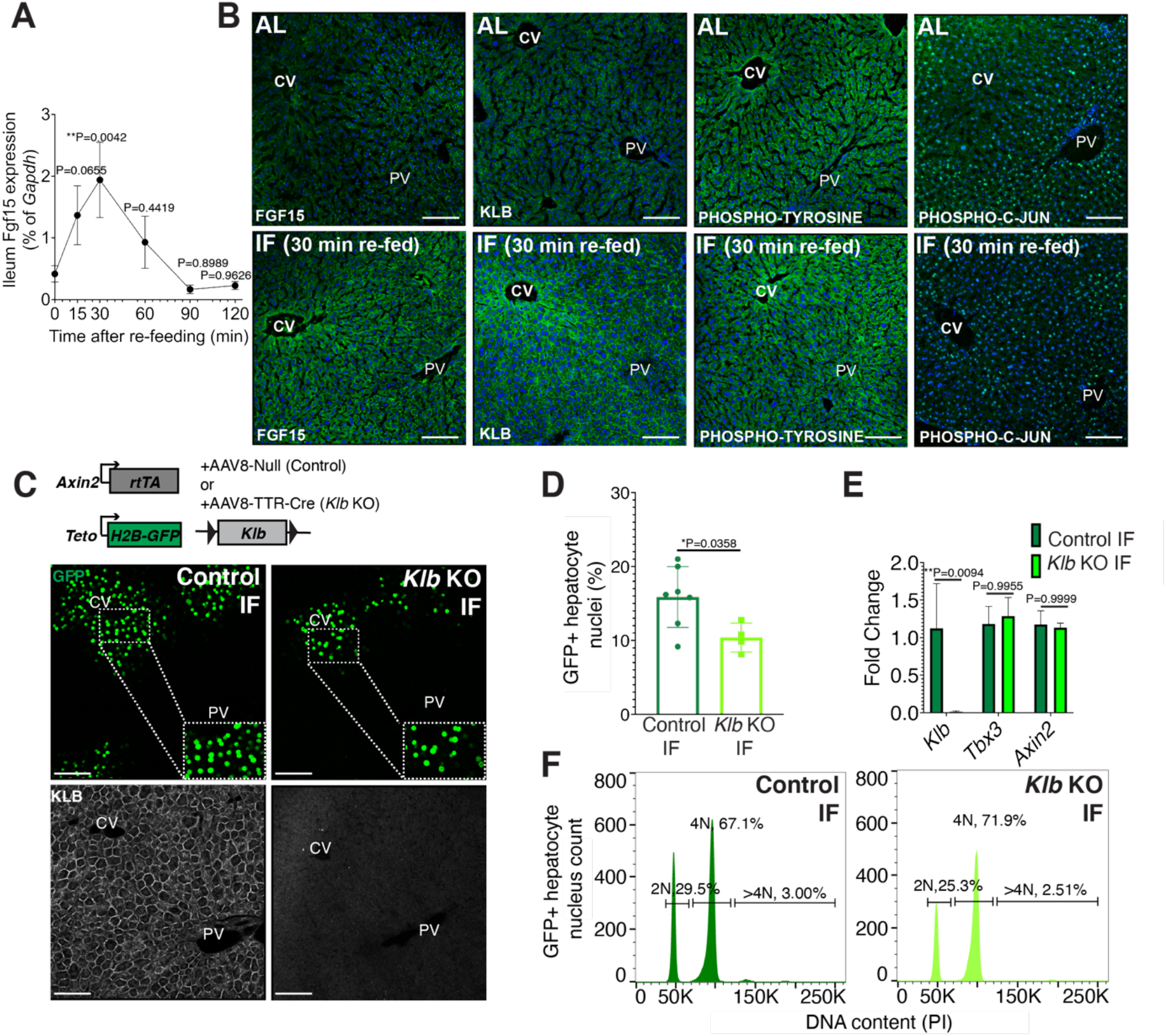
Endocrine FGF15-ß-KLOTHO(KLB) signaling is required for hepatocyte proliferation during intermittent fasting. **(A)** Quantitative real-time PCR analysis hightlighing rapid increase in *Fgf15* expression in ileum 30mins after re-feeding in 1-week IF-treated livers. One-way ANOVA, comparison with time 0, N=3. **(B)** Immunofluorescence for endocrine FGF pathway components highlighting pathway activation in 1 week IF-treated livers 30 mins after re-feeding, (ZT12).**(C)** Schematic of method to deplete hepatocytes of *Klb*. *Axin2-rtTA*; *Teto-H2BGFP*; *Klbflox/flox* mice were injected with AAV8-TTR-Cre (*Klb* KO). GFP and KLOTHO immunofluorescent images showing decrease in hepatocyte expansion and loss of KLOTHO in *Klb* KO compared to Control livers. **(D)** Percentage of GFP+ hepatocyte nuclei in *Klb* KO and Control livers. **(E)** Quantitative real-time PCR analysis confirming loss of *Klb* but not Wnt target genes, *Tbx3* and *Axin2*, in *Klb* KO livers. 2-way ANOVA, N=3. **(F)** Ploidy distribution of GFP+ hepatocyte nuclei in Control IF and *Klb* KO IF livers. Unpaired t-test. N= 7 (Control), 4 (*Klb* KO). \*\**P*<0.01, \**P*<0.05.Error bars indicate standard deviation.Scale bar, 100μm.

Given that endocrine FGF15 is an early regulator of the hepatocyte response to food intake, we asked whether loss of endocrine FGF signaling in hepatocytes would prevent IF-induced proliferation. To test this, we genetically depleted hepatocytes of the endocrine FGF receptor (*Klb*) and traced expansion of Axin2+ GFP-labeled cells during IF-treatment (Figure 2C). Loss of *Klb* led to a 53% reduction in GFP-labeled pericentral hepatocyte nuclei compared to control animals after 1 week of an IF feeding regimen (Figure 2D). Importantly, loss of *Klb* did not impact WNT target gene expression (Figure 2E) a critical regulator of hepatocyte zonation and function in the liver (Perugorria et al., 2019; Wang et al., 2015). Furthermore, loss of *Klb* did not significantly change GFP-labeled nuclei ploidy (Figure 2F). In summary, these data emphasize the functional requirement for endocrine FGF15-β-KLOTHO signaling to promote IF-induced pericentral hepatocyte proliferation.

### WNT signaling and the WNT target gene *Tbx3* promote enhanced pericentral hepatocyte proliferation during intermittent fasting

Next, we sought to understand why pericentral hepatocytes preferentially divided in response to IF, as FGF15 is a hormone and would be theoretically accessible to all hepatocytes. One hypothesis is that IF-induced hepatocyte proliferation additionally requires a second signal, which is concentrated near pericentral hepatocytes. Pericentral hepatocytes receive paracrine WNT signals from endothelial cells of the central vein, which is required to establish and maintain pericentral hepatocyte zonation and function in the liver (Perugorria et al., 2019; Wang et al., 2015). We tested whether ectopic and constitutive activation of the WNT pathway in midlobular and periportal hepatocytes, which typically do not receive WNT signaling, would promote hepatocyte proliferation under an IF feeding regimen. To ectopically activate WNT signaling, we genetically deleted the WNT repressor *Apc* by injecting mice ubiquitously expressing *Cas9* with an AAV8 carrying a guide RNA (sgRNA) directed against *Apc* (Figure 3A). Mid-lobular and periportal hepatocytes that had undergone CRISPR-Cas9 gene editing of *Apc*, were identified by the expression of WNT target gene glutamine synthetase (GS). Remarkably, these WNT activated (GS+) cells remained mostly as single cells in AL treatment, however they clonally expanded under IF treatment (Figure 3A and B).

**Figure 3.**
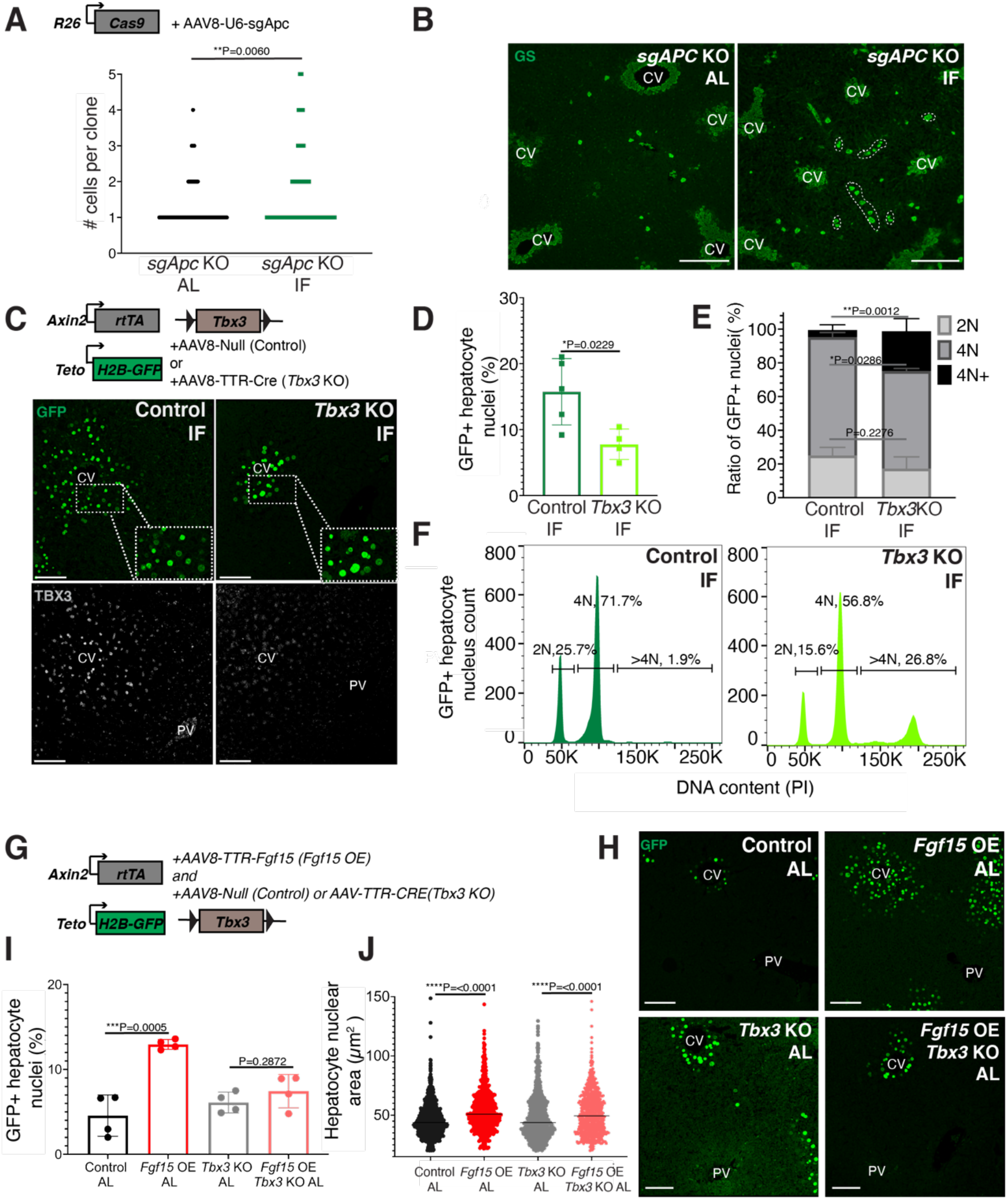
Paracrine WNT and WNT target gene *Tbx3* promotes hepatocyte proliferation during intermittent fasting. **(A)** Method to constitutively activate WNT signaling in midlobular, periportal cells. AAV8-U6-sgAPC was injected into the tail vein of *R26-Cas9* mice. Animals were IF-treated for 1 week before analysis. GS immunofluorescent images for detection of *Apc* mutant clones in AL and IF livers. **(B)** Number of *Apc* mutant hepatocytes per 3D clone expand in IF compared to AL livers. Mann-Whitney test. 130 clones analyzed in AL. 74 clones analyzed in IF. N=3. **(C)** Schematic of method to deplete hepatocytes of the WNT target, *Tbx3*. *Axin2-rtTA*; *Teto-H2BGFP*; *Tbx3flox/flox* mice were intraperitoneally injected with AAV8-TTR-Cre (Tbx3 KO). GFP and TBX3 immunofluorescent images to show IF-induced proliferation and *Tbx3* depletion, respectively, in Control and *Tbx3* KO livers. **(D)** Percentage of GFP+ hepatocyte nuclei decreased in *Tbx3* KO IF compared to Control IF livers. Unpaired t-test, N=7(Control IF), N=4(Tbx3 KO IF). **(E** and **F)** Nuclear ploidy distribution of GFP+ hepatocytes highlighting hyper-polyplodization in *Tbx3* KO IF compared to Control IF livers. Two-way ANOVA, N= 3. **(G)** Schematic for *Fgf15* overexpression. *Axin2-rtTA*; *Teto-H2BGFP*; *Tbx3 flox/flox* mice were injected with AAV-TTR-FGF15(*Fgf15* OE) and AAV8-Null(Control) or AAV-TTR-CRE(*Tbx3* KO). **(H)** GFP immunofluorescent images from Control AL, *Fgf15* OE AL, *Tbx3* KO AL, *Fgf15* OE; *Tbx3* KO AL livers. **(I)** Percentage of GFP+ hepatocyte nuclei highlighting lack of hepatocyte proliferation in *Tbx3* KO livers. Unpaired t-test, N=4. **(J)** Dot plot highligting increase in nuclear area with *Fgf15* overexpression both with and without *Tbx3*. Unpaired t-test. \*\*\*\**P*<0.0001, \*\*\**P*<0.001, \*\**P*<0.01, \**P*<0.05. Error bars indicate standard deviation. Scale bar, 100μm.

WNT signaling induces expression of the transcriptional repressor *Tbx3* in hepatocytes. Previous studies have demonstrated that *Tbx3* regulates division of hepatocytes during liver development by repressing cell cycle inhibitors(Jin et al., 2022; Suzuki, Sekiya, Buscher, Izpisua Belmonte, & Taniguchi, 2008). To determine if *Tbx3* plays a role in IF-induced hepatocyte proliferation, we genetically depleted *Tbx3* in hepatocytes and traced expansion of Axin2+ GFP-labeled cells after IF-treatment (Figure 3C). Loss of *Tbx3* led to a 51% reduction in expansion of GFP-labeled cells after IF-treatment (Figure 3D). Furthermore, loss of *Tbx3* increased nuclear ploidy in GFP-labeled cells, suggesting that pericentral hepatocytes underwent endoreplication rather than division (Figure 3E and F). These findings suggest that WNT and WNT-induced transcription factor TBX3 endow pericentral hepatocytes with capacity to divide during IF treatment.

### FGF15 signaling requires WNT/TBX3 to induce pericentral hepatocyte proliferation

Our results suggest that systemic FGF15 and paracrine WNT pathways may work together to push hepatocytes through the cell cycle. To directly test for an interdependent relationship between FGF and WNT signaling on hepatocyte division, we ectopically expressed *Fgf15* in the liver under AL feeding in the presence or absence of *Tbx3* (Fig. 3 G, H). Indeed, AAV-mediated *Fgf15* overexpression, led to a 102% percent increase in Axin2+ GFP-labeled nuclei compared to control animals, (Figure 3I). However, *Fgf15* overexpression with *Tbx3* loss, did not significantly increase hepatocyte division (Figure 3I). Interestingly, *Fgf15* overexpression increased nuclear area both with and without loss of *Tbx3* (Figure 3J), suggesting that FGF signaling initiates S-phase, but requires WNT through TBX3 to complete mitosis. These results highlight the co-requirement of FGF15 and WNT signaling for pericentral hepatocyte proliferation.

### Hepatocyte proliferation or compensatory polyploidization maintains the hepatostat during intermittent fasting

During partial hepatectomy where two thirds of the liver is removed or during liver transplantation from a smaller organism to a larger one, loss of liver cell mass disrupts the hepatostat, the liver-to-body-weight ratio required to maintain homeostasis (Michalopoulos & Bhushan, 2021). Re-establishment of the hepatostat through hepatocyte regeneration is critical to prevent development of liver disease (Michalopoulos & Bhushan, 2021). We asked whether the hepatostat was disrupted during intermittent fasting. For early timeframes of IF (2-6 days), liver-to-body weight ratio significantly decreased during fasting and increased during re-feeding states compared to AL-treated livers (Figure 4A). However, after 3 weeks of IF-treatment, this ratio stabilized and was not significantly different between fasting, re-feeding or AL states.

**Figure 4.**
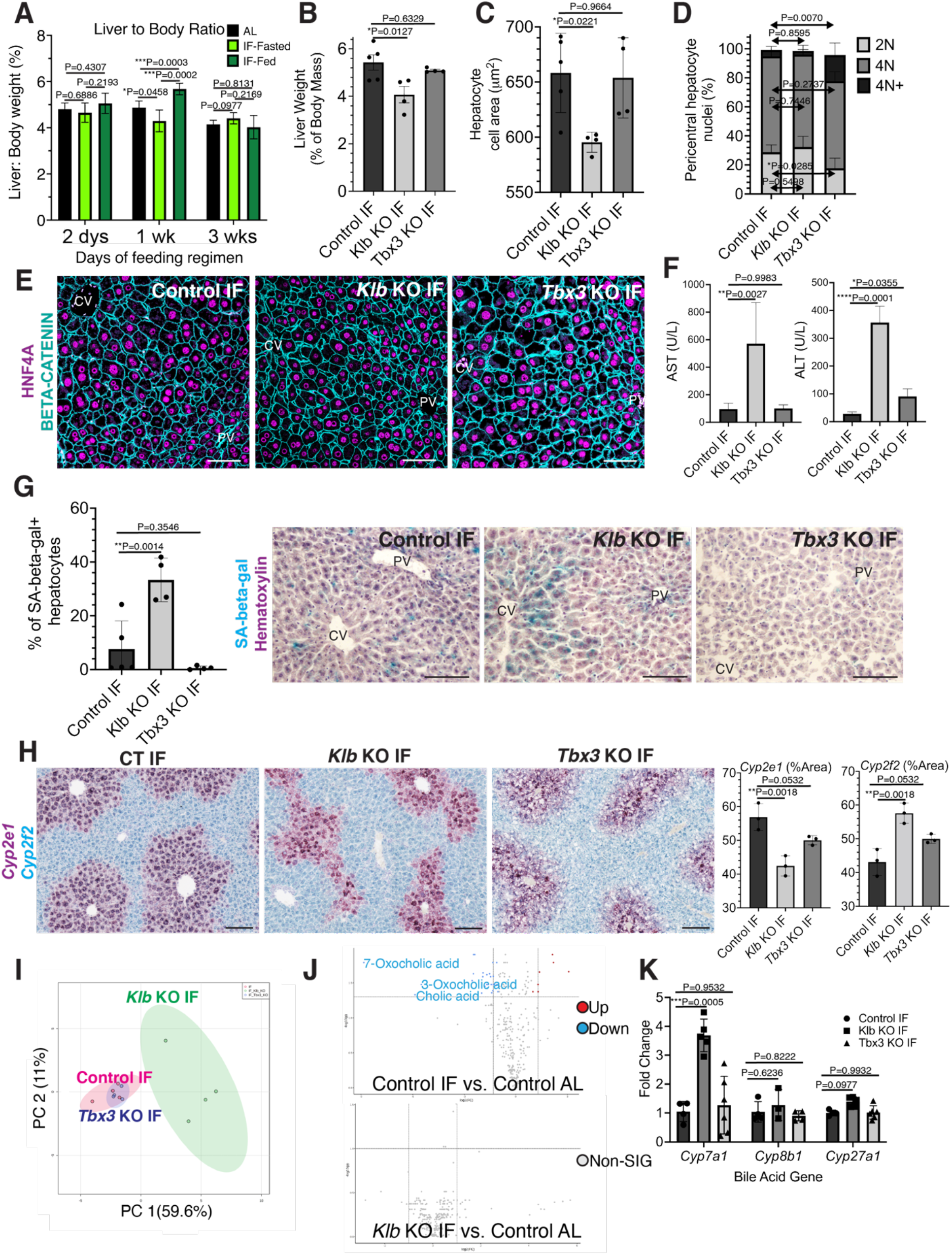
Hepatocyte proliferation or compensatory polyploidization maintains the hepatostat during intermittent fasting. **(A)** Liver to body weight ratio in wild type livers during 2 dy, 1 week and 3 weeks of IF and AL feeding. **B-K** Liver analyses after 3 weeks of IF treatment in Control, *Klb* KO and *Tbx3* KO livers. **(B)** Liver to body weight ratio. **(C)** Hepatoycte nuclear area. **(D)** Nuclear ploidy distribution of pericentral hepatocytes with hyper-polyplodization in *Tbx3* KO IF livers. **(E)** Immunofluorescenceimages for BETA-CATENIN and HNF4A highlighting hepatocyte cell and nuclear areaduring IF. **(F)**AST and ALT liver injury marker presence in serum. **(G)** Quantification and representative images of senescence-associated beta-galatosidase stains. **(H)** RNAscope images and quantification for pericentral marker Cyp2el and periportal marker Cyp2f2. **(I)** Metabolomics PCA plot comparing Control IF, *Klb* KO IF and *Tbx3* KO IF livers. **(J)** Volcano plots comparing metabolites between Control AL and Control IF livers and Control AL and *Klb* KO IF livers. The top 3 most significantly changed bile metabolites are labeled in blue. **(K)** Expression of bile acid pathway enzymes genes in livers. Quantitative PCR for genes for critical enzymes in bile acid pathway. All statistics were performed on 3-5 animals using 1-way ANOVA. \*\*\*\**P*<0.0001, \*\*\**P*<0.001, \*\**P*<0.01, \**P*<0.05. Error bars indicate standard deviation. Scale bar, 100μm.

Given the reestablishment of the hepatostat after 3 weeks of intermittent fasting, we then asked if the transient hepatocyte proliferation observed during early timeframes of IF was required to reach homeostasis after 3 weeks of IF. To test this, we compared the liver-to-body weight ratios, hepatocyte cell and nuclear areas between livers depleted of *Klb* or *Tbx3* and ad libitum fed or intermittently fasted for 3 weeks. In AL-treated animals, loss of *Klb* did not significantly alter the liver-to-body weight ratio nor hepatocyte cell and nuclear area compared to control livers (Supplementary Figure 3 A-D). In contrast, in IF-treated animals, loss of *Klb* led to a significant decrease compared to control livers (Figure 4B-E). Livers depleted of *Tbx3* were able to maintain liver-to-body weight ratios, hepatocyte cell and nuclear area similar to control livers during AL and IF-treatment (Figure 4B-E, Supplementary Figure 3 A-D). Importantly, IF-treated livers depleted of *Tbx3* exhibited hyper polyploidization of pericentral hepatocytes (Figure 4D). These findings highlight the importance of hepatocyte proliferation during IF to maintain the hepatostat. Moreover, they demonstrate the ability of hepatocytes to undergo endoreplication and polyploidization, in the absence of division, as a compensatory mechanism to maintain liver size.

To determine the functional consequence of loss of hepatocyte proliferation during IF, we analyzed fibrosis and cell death in control, *Klb* KO, or *Tbx3* KO livers under IF or AL feeding regimens (Supplementary Figure 4 A-C). Despite the significant reduction in liver-to-body weight ratio with loss of *Klb* during IF, we did not observe an increase in fibrosis or dying (TUNEL positive) hepatocytes with loss of *Klb* or loss of *Tbx3* during IF-treatment. However, we did observe a greater than 3-fold increase in transaminase AST and ALT—widely used markers of liver injury—in serum from *Klb* KO IF-treated livers compared to control IF and *Tbx3* KO IF-treated livers (Figure 4F). We also observed a striking increase in pericentral-localized hepatocyte senescence in *Klb* KO IF-treated livers compared to control IF and *Tbx3* KO IF-treated livers (Figure 4G). Furthermore, hepatocyte senescence and serum transaminase levels were elevated by 2-fold in *Klb* KO IF-treated livers compared to *Klb* KO AL treated livers (Figure 4F, G and Supplementary Figure 3 E,F). These two findings demonstrate that when compensatory hepatocyte proliferation is blocked during IF, liver pathology ensues.

To further support these findings, we performed zonation studies and metabolomics to identify additional functional consequences of compromised hepatocyte proliferation during intermittent fasting. These analyses highlighted marked changes in zonation (Figure 4H) as well as liver metabolites (Figure 4I) between control IF and *Klb* KO IF treated livers. Curiously, differences in metabolites between IF and AL samples were lost when *Klb* was lost during IF treatment (Figure 4J, Supplementary Table 1) indicating that the hepatocyte metabolism required to maintain the hepatostat during IF was impaired when hepatocyte proliferation and polyploidization was lost. To further test this, we examined the zonation and expression of genes important in production of bile acids, the most strongly changed metabolites between IF and AL. Importantly, Cyp7a1, a gene that is pericentrally expressed in the normal liver(Halpern et al., 2017), is dysregulated by 3-fold with loss of *Klb* during IF treatment (Figure 4K). These data combined demonstrate that loss of division of pericentral cells during IF has important functional consequences for the liver including a decrease in the liver-to-body-weight ratio, increased pericentral hepatocyte senescence and irregular liver zonation and metabolism.

## Discussion

The liver is thought to be mostly quiescent except in the presence of injury. Our data overturn this view by demonstrating that the liver is exquisitely tuned to changes in nutrient status and deploys robust homeostatic mechanisms to ensure a constant liver to body weight ratio during fasting and re-feeding. In order to proliferate in response to intermittent fasting, hepatocytes integrate nutrient sensing responses, mediated via intestinal FGF15, with knowledge of cellular position, mediated by local, pericentral WNT signals. Pericentral hepatocyte proliferation ensures replacement of lost cellular mass through increases in overall cell numbers. We propose a working model in which paracrine WNT and endocrine FGF pathways work together to push hepatocytes through different phases of the cell cycle. The role of FGF15 in pericentral hepatocyte proliferation observed in our studies suggest that this signal initiates S-phase, whereas WNT through TBX3 permits progression through M-phase. It has been demonstrated in other contexts as well that both FGF and WNT signaling are conjointly required for tissue growth (McGrew, Hoppler, & Moon, 1997; ten Berge, Brugmann, Helms, & Nusse, 2008).

It will be of interest to know how the hepatostat is maintained during other nutrient conditions such as ketogenic, calorie restricted, or high-fat diets. Many tissues in the body are known to be slowly proliferating, but this conclusion has mainly stemmed from AL-fed mouse studies, which may not mimic the intermittent fasting that wild animals are exposed to during times of fluctuating food abundance. These studies demonstrate that there is perhaps more proliferative capacity than previously appreciated in tissues that are currently thought to exhibit a slow turnover rate. This would have implications for how we understand regulation of both stem and non-stem cell populations in many other tissues and organs.

## Materials and Methods

### Key Resource Table

| Antibody Target | Company (Catalogue Number) RRID | Dilution |
| --- | --- | --- |
| GFP | Abcam Cat# ab13970<br>RRID:AB_300798 | 1:500 IF |
| RFP | Rockland Cat# 600-401-379<br>RRID:AB_2209751 | 1:500 IF |
| HNF4-ALPHA | Abcam Cat# ab41898<br>RRID:AB_732976 | 1:500 IF; 50 IHC |
| KLOTHO | DSHB Cat# Klotho KL-115<br>RRID:AB_2618099 | 1:50 IF |
| FGF15 | Santa Cruz sc-514647<br>RRID NA | 1:50 IF |
| PHOSPHO-TYROSINE | Cell Signaling Technology Cat# 9411<br>RRID:AB_331228 | 1:50 IF |
| PHOSPHO-C-JUN | Cell Signaling Technology Cat# 3270,<br>RRID:AB_2129575 | 1:50 IF |
| TBX3 | Santa Cruz Biotechnology Cat# sc-17871<br>RRID:AB_661666 | 1:50 IHC |
| GOAT IgG | Jackson ImmunoResearch Labs Cat# 705-065-147<br>RRID:AB_2340397 | 1:200 IHC |
| GLUTAMINE SYNTHETASE | Millipore Cat# MAB302<br>RRID:AB_2110656 | 1:500 IHC |
| BETA-CATENIN | BD Biosciences Cat# 610154<br>RRID:AB_397555 | 1:50 IHC |
| HNF4-ALPHA | Santa Cruz Biotechnology Cat# sc-8987<br>RRID:AB_2116913 | 1:50 IHC |
| KI67 | Thermo Fisher Scientific Cat# 14-5698-82<br>RRID:AB_10854564 | 1:50 IHC |

| Mice Used | Company (Strain) RRID |
| --- | --- |
| *C57BL/6J* | Jax(000664) RRID:IMSR_JAX:000664 |
| *R26-CreERT2* | Jax (008463) RRID:IMSR_JAX:008463 |
| *R26-Confetti* | Jax (017492) RRID:IMSR_JAX:008463 |
| *Axin2-rtTA* | Jax (016997) RRID:IMSR_JAX:016997 |
| *TetO-H2B-GFP* | Jax (005104) RRID:IMSR_JAX: 005104 |
| *R26-mTmG* | Jax (037456) RRID:IMSR_JAX: 037456 |
| *R26-Cas9* | Jax (026179) RRID:IMSR_JAX: 026179 |

### Mouse strains, husbandry, and experimental methods

All experiments were done on adult, 8–12-week-old, male mice, unless otherwise noted. Wild type *C57BL/6J* mice, *R26-CreERT2(Ventura et al., 2007), R26-Confetti (Snippert et al., 2010), Axin2-rtTA(Yu et al., 2007), TetO-H2B-GFP(Tumbar et al., 2004);R26-mTmG(Muzumdar, Tasic, Miyamichi, Li, & Luo, 2007)* and *R26-Cas9(Platt et al., 2014)* strains were obtained from The Jackson Laboratory (JAX, Bar Harbor, ME, see Key Resources Table). *Tbx3* flox mice were a gift from Dr. Anne Moon(Frank et al., 2012). *Klb* flox mice were a gift from Dr. David Mangelsdorf(Ding et al., 2012). All mice were housed in the animal facility of Stanford University on a 12-h light/dark cycle (0700h/1900h) ad libitum access to water and food (standard chow diet with 18% calories derived from fat; 24% calories from protein and 58% calories from carbohydrates, Tekland 2918).

For all intermittent fasting, mice were randomly assigned into ad libitum or intermittent fasting groups. Intermittent fasting was performed with total food deprivation and ad libitum access to water from approximately 1900 h to 1900 h the following day to implement alternate periods of 24 hour fasting and feeding. Unless specified otherwise in text and figures, all samples were collected for both ad libitum and intermittent fasting groups at 1200 h to access samples during a neutral metabolic and circadian rhythm time point.

For random cell lineage tracing studies, *R26-CreERT2*/*R26-Conf*etti mice received intraperitoneal injections of tamoxifen (TAM; 4mg/25 grams mouse weight, Sigma, St. Louis, MO) dissolved in 10% ethanol/corn oil (Sigma, St. Louis, MO) twice with 48 hours between injections. 2 weeks after the last tamoxifen injection, livers where immediately analyzed (T0) or analyzed after an additional 1-3 weeks of ad libitum feeding or intermittent fasting.

For Axin2+ cell lineage tracing studies, mice received doxycycline hycalate (Dox; 1mg/ml; Sigma, St. Louis, MO) in drinking water for 5 days. Dox water was then replaced with normal drinking water for 3 days before livers were immediately analyzed(T0) or analyzed after an additional 1 week, 3 weeks and 3 months of ad libitum feeding or intermittent fasting

For Cyp1a2-CreER lineage tracing studies, mice received a single dose of 100mg/kg body weight of tamoxifen intraperitoneally. 2 weeks after the last tamoxifen injection, livers where immediately analyzed (T0) or analyzed after an additional 1-3 weeks of ad libitum feeding or intermittent fasting.

For all AAV studies, mice were intraperitoneally injected with 1×10^11^ genome copies per mouse at 6-8 weeks of age to induce liver specific depletion of *Klb* or *Tbx3(AAV8-TTR-Cre*, Vector Bio Labs, Malvern, PA), *Fgf15* overexpression (AAV8-TTR-FGF15) or *Apc* gene editing (AAV8-U6-sgAPC). For AAV control studies, an AAV8-Null vector containing no transgene (Vector Bio Labs, Malvern PA) was used on a combined cohort of *Tbx3 flox/flox* and *Klb flox/flox* mice. For studies that combined AAV injection and Axin2+ cell lineage tracing, mice were first injected with AAV, allowed 3 days to recover and subsequently treated with dox water to induce tracing.

All animal experiments and methods were approved by the Institutional Animal Care and Use committee at Stanford University. In conducting research using animals, the investigators adhered to the laws of the United States and regulations of the Department of Agriculture.

### Tissue collection, processing, staining, and imaging

For clonal analysis and KLOTHO immunofluorescence, mice were perfused with 4% paraformaldehyde (PFA), livers were isolated and further fixed in 4% PFA for 2 hours at 4°C. PFA-fixed tissues were washed in PBS and sectioned into 50-200μm sections using a Compresstome vibrating microtome tissue slicer (VF-310-0Z, Precisionary). Vibratome sections from the median lobe where then permeabilized, stained and cleared using a method developed by the Zerial lab (www.zeriallab.org). Antibodies and dilutions described in Key Resources Table. Vibratome sections were imaged using an SP8 White Light Laser Confocal microscope (Lecia; Weltzar, Germany) and a BZ-X800 microscope (Keyence; Osaka, Japan). Confocal image stacks were acquired at 20x magnification and up to 100 μm with a step size of 1 μm along the Z-axis and processed and analyzed with Imaris software.

For endocrine FGF signaling pathway analyses (Figure 2B), livers were flash-frozen in OCT, cryo-sectioned at 10μm, fixed for 15 mins in 4% PFA at room temperature and then stained. Antibodies and dilutions described in Key Resources Table.

For histology and immunohistochemistry, liver was fixed overnight in 10% formalin at room temperature, dehydrated, cleared in HistoClear (Natural Diagnostics), and embedded in paraffin. Sections were cut at 5μm thickness, de-paraffinized, re-hydrated and processed for further staining via immunofluorescence or *in situ* hybridization assays as described below.

For histology, formalin-fixed paraffin-embedded liver sections were sent to the Department of Comparative Medicine’s Animal Histology Services for Sirius Red staining.

For immunofluorescence, sections of formalin-fixed paraffin-embedded livers were subjected to antigen retrieval with Tris buffer PH=8.0 (Vector Labs H-3301; Newark CA) in a pressure cooker. They were then blocked in 5% normal donkey serum in PBS containing 0.1% Triton-X, in combination with the Avidin/Biotin Blocking reagent (Vector Labs SP-2001; Newark CA). Sections were incubated with primary and secondary antibodies and mounted in Prolong Gold with DAPI medium (Invitrogen; Waltham, MA). Biotinylated goat antibody was applied to section stained with TBX3, before detection with Streptavidin-647. Antibodies and dilutions described in Key Resources Table. Samples were imaged at 20X magnification using an Sp8 Confocal or a Zeiss Imager Z.2 and processed and analyzed with ImageJ software.

### Senescence-associated Beta-galactosidase staining and TUNEL assay

For senescence-associated beta-galactosidase staining, flash frozen livers were cryosectioned at 10μm and fixed with 0.5% glutaraldehyde in PBS for 15 mins, washed with PBS supplemented with 1mM MgCl_2_ and stained for 14 hrs in PBS containing (1 mM MgCl2.; 1 mg/ml X-Gal and 5 mM of each of Potassium ferricyanide and Potassium ferrocyanide). Assay was performed at pH= 5.5 as previously described (Krizhanovsky et al., 2008). Hematoxylin was used as a counterstain.

To detect apoptosis in livers, formaldehyde-fixed paraffin-embedded sections were detected by terminal deoxynucleotidyl transferase-mediated dUTP-biotin nick end labeling (TUNEL) assay (ThermoFisher Scientific; C10619; Waltham, MA) according to the manufacturer’s instructions.

### RNAscope *in situ* hybridization

*In situs* were performed using the RNAscope 2.5 HD Duplex Reagent Kit (Advanced Cell Diagnostics; Newark, CA) according to the manufacturer’s instructions. Images were taken at 20x magnification on a Zeiss Imager Z.2 and processed using ImageJ software. Probes used in this study were *Cyp2f2* (target region: 555-1693) and *Cyp2e1* (target region: 458-1530).

### Fibrosis Assay

Bright-field images were collected on a Zeiss Imager Z.2. Red stained collagen levels were quantified using image J (imagej.nih.gov).

### Clone size, number, and location

To quantify clone size, threshold of fluorescent channels was lowered so that clear cell and nuclear boundaries could be distinguished. 1 large-stitched image with an area of 1.8 x 1.8 x 0.1 mM^3^ mouse was taken from 2 representative vibratome sections from each mouse at each time point. Only clones completely within the tissue sample were analyzed. We counted the total number of clones from the 6 representative images. Because images were of equal area, clone numbers can be compared to each other between time points. Clones containing a cell located within 3 cell distances from the portal vein or bile duct were classified as periportal; clones containing a cell within 3 cell distances from the central vein were classified as pericentral; clones not meeting either criterion were classified as midlobular.

### Hepatocyte nuclei isolation and analysis

For hepatocyte nuclei isolation, liver lobes from mice were homogenized in cold 1% formaldehyde in PBS with a loose pestle and Dounce homogenizer. Samples where then fixed for 10 min at room temperature followed by incubation for 5 min with glycine at a final concentration of 0.125 M. Samples were centrifuged at 300 g for 10 min, at 4°C. Pellets were washed in PBS and re-suspended with 10 ml cell lysis buffer (10 mM Tris-HCL, 10 mM NaCl, 0.5% IGEPAL) and filtered through 100 μm cell strainers. A second round of homogenization was performed by 15-20 strokes with a tight pestle. Nuclei were pelleted at 2000 g for 10 min at 4°C and re-suspended in 0.5ml PBS and 4.5 ml of pre-chilled 70% ethanol and stored at −20°C before downstream GFP content and ploidy analysis by flow cytometry.

Right before flow cytometry, 1 million nuclei were resuspended in PBS and stained with FxCycle PI/RNase (Thermo Fisher, F10797; Waltham, MA) staining solution for 15-30 minutes at room temperature. Cells were analyzed on a FACS ARIA II (BD). Data were processed with FACS Diva 8.0 software (BD) and FlowJo v10 (FlowJo). Doublets were excluded by FSC-W×FSC-H and SSC-W×SSC-H analysis. Single-stained channels were used for compensation and fluorophore minus one control was used for gating.

### Real-time PCR measurement

Liver samples were homogenized in TRIzol (Invitrogen; Waltham, MA) with a bead homogenizer (Sigma, St. Louis, MO). Total RNA was purified using the RNeasy Mini Isolation Kit (Qiagen, Hilden, Germany) and reverse-transcribed (High Capacity cDNA Reverse Transcription Kit; Life Technologies, Carlsbad, CA) according to the manufacturer’s protocol. Quantitative RT-PCR were performed with TaqMan Gene Expression Assays (Applied Biosystem, Waltham, MA) on a StepOnePlus Real-Time PCR System (Applied Biosystems, Waltham, MA). Relative target gene expression levels were calculated using the delta-delta CT method(Livak & Schmittgen, 2001). Gene Expression Assays used were *Gapdh* (Mm99999915_g1) as control, *Klb* (Mm00473122_m1), *Axin2* (Mm00443610_m1), *Fgf15*(Mm00433278_m1), *Tbx3* (Mm01195719_m1), *Cdnk1a(Mm00432448)* all from Thermo Fisher Scientific (Waltham, MA).

### Single Cell RNA Sequencing

Hepatocytes were isolated from livers of 8-week-old C57BL/6J mice that had been intermittent fasted for 1 week or ad libitum fed using a two-step collagenase perfusion technique as previously described(Peng et al., 2018).

Collections were performed at 1000h and during the feeding cycle of IF. For each feeding regimen, 3 livers were collected and processed as 3 individual samples. For each sample, 2,000 hepatocytes were loaded to target ~1000 cells after recovery according to the manufacturer’s protocol. Single cell libraries were prepared using the 10x Genomics Chromium Single Cell 3” Reagents Kit V3. Single cell libraries were loaded on an Illumina NovaSeq 6000 instrument with NovaSeq S2 v.1.5 Reagent Kits with the following reads: 28 bases Read 1 (cell barcode and unique molecular identifier (UMI)), 8 bases i7 Index 1 (sample index), 91 bases Read 2 (transcript).

Sample demultiplexing, barcode processing, single-cell counting and reference genome mapping were performed using the Cell Ranger Software (v3.1.0, mm10 ref genome) accordingly to the manual. All samples were normalized to present the same effective sequencing depth by using Cell Ranger aggr function. The dimensionality reduction by principal components analysis (PCA), the graph-based clustering and UMAP visualization were performed using Seurat (v3.0, R package). Genes that were detected in less than three cells were filtered out, and cells were filtered out with greater than 10 percent of mitochondrial genes and with fewer than 200 or greater than 50000 detected genes.

For cell clustering, R software was used to sort cells into either pericentral (PC), midlobular (Mid), and periportal (PP) classes based on the greatest expression of biomarkers *Cyp2e1*, *Cyp1a2*, *Glul* (for PC), *Hamp* and *Igfbp2* (for Mid), *Cyp2f2* and *Cps1* (for PP).

### Generation of AAV Expression Vectors

The AAV-TTR-FGF15 virus was produced from the complementary stand AAVS construct, csAAV-TTR-CRE plasmid (kind gift of Holger Willenbring). The *CRE* gene was excised by digestion with SalI (NEB). The *Fgf15* gene (GenBank: BC021328 cloneID 5066286) was amplified for assembly into the SalI cut AAV-TTR backbone with NEB HiFi Builder using the primers: (FWD) 5’ggagaagcccagctgGTCGACGCCACCATGGCGAGAAAGTGGAACGG 3’ and (REV) 5’ atcagcgagctctaGTCGACTCATTTCTGGAAGCTGGGACTCTTCAC 3’. The two fragments were assembled with NEB HiFi builder and cloned in NEB Stable e. coli.

The AAV-sgApc virus was produced from the pAAV-Guide-it-Down construct (Clontech Laboratories Inc., 041315) using assembly primers:

(FWD) 5’CCGGAGGCTGCATGAGAGCACTTG3’ and
(Rev) 5’AAACCAAGTGCTCTCATGCAGCCT3’3.
AAV-sgApc contains a U6 promoter and an sgRNA targeting the sequence
5’AGGCTGCATGAGAGCACTTG3’ in exon 13 of *Apc*.

### Metabolite Extraction

Livers were harvested and immediately flash frozen in LN2 then stored at −80°C. While kept on dry ice a 20 mg sample was removed from each liver specimen, massed using an analytical balance, and placed in a 2 mL round bottom polypropylene tube containing 4-6, 2.3 mm stainless steel beads. 500 μL of −20°C extraction solution (methanol: acetonitrile: water, 2:2:1) containing stable isotope labeled metabolite standards was added to each sample tube. Ratio of 20 mg to 500 μL was retained when masses were not exactly 20 mg. All samples were homogenized at an amplitude of 20 Hz for 15 minutes and stored at −20°C for one hour to maximize protein precipitation. Samples were then vortexed for 20 seconds and centrifuged at 4°C for 5 minutes, speed 14,000 rcf. 120 μL of supernatant was removed from each tube and filtered using 0.2 μm polyvinylidene fluoride filter (Agilent Technologies P/N: 203980-100) and collected via 6,000 rcf centrifuge for 4 minutes. An additional 50uL was removed from each sample and combined into 5 pooled samples analyzed at equal intervals throughout the analysis to ensure stable signal. Extracts, pools, and procedural blanks were sealed and stored at 4°C until prompt analysis.

### HILIC-MS/MS Metabolite Data Collection and Processing

Untargeted metabolomics analysis was conducted as described previously(Han et al., 2021) with some modification. Liver extracts were analyzed via hydrophilic interaction liquid chromatography (HILIC) coupled to a Thermo Q-Exactive HF high resolution mass spectrometer. Each sample was analyzed in both positive and negative ionization modes (ESI+, ESI-) via subsequent injections. Full MS-ddMS2 data was collected, an inclusion list was used to prioritize MS2 selection of metabolites from our in-house ‘local’ library, when additional scan bandwidth was available MS2 was collected in a data-dependent manner. Mass range was 60-900 mz, resolution was 60k (MS1) and 15k (MS2), centroid data was collected, loop count was 4, isolation window was 1.5 Da. Metabolomics data was processed using MS-DIAL v4.60 (Tsugawa et al., 2020) and queried against a combination of our in-house MS2 library(Han et al., 2021) and MassBank of North America, the largest freely available spectral repository(Kind et al., 2018). Annotations were scored using guidelines from the metabolomics standards initiative(Members et al., 2007). Features were excluded from analysis if peak height was not at least 5-fold greater in one or more samples compared to the procedural blank average. Statistical analysis of annotated features was implemented using MetaboAnalyst 5.0(Pang et al., 2021). Data visualization including principal component analysis and volcano plots were generated using log10 transformed peak heights.

## Data Availability

Single Cell RNA Sequencing will be made available on Gene Expression Omnibus(GEO) after approval by GEO.

Metabolomics will be made available on Metabolomics Workbench after approval by NIH Common Fund’s National Metabolomics Data Repository (NMDR).

## Acknowledgments

We thank K. M. Loh and Natalie Torok for constructive feedback on the manuscript; the Chan Zuckerberg Biohub Community Access Program for use of 10x Genomics equipment and sequencing; Stanford Neuroscience Gene and Vector Virus Core for virus production; and H. Willenbring for sharing the csAAV-TTR-CRE plasmid. This study was supported by the Howard Hughes Medical Institute (HHMI) and the Stinehart Reed Foundation. A.S. was supported by the Office of the Assistant Secretary of Defense for Health Affairs, through the Peer Reviewed Cancer Research Program, under Award No. W81XWH-17-1-0245. Opinions, interpretations, conclusions, and recommendations are those of the author and are not necessarily endorsed by the Department of Defense. P.W. was supported by the Damon Runyon Cancer Research Foundation (DRSG-28P-19). The Genetics Bioinformatics Service Center of Stanford Cancer Institute’s Share Resource Facility was supported by NIH grant P30 CA124435.

## Additional Information

A.S. and R.N. conceived the project, performed experiments, and wrote the manuscript. Y. J. performed flow cytometry and analyses. B. D. performed metabolomics and analyses. C.L. performed TUNEL analyses. P.W., T.A., C.L., M.M., and N.N. helped generated scRNA-seq data. Y.Y. helped processed and analyzed scRNA-seq data. H.N. helped performed animal experiments. T.A. M.F., and A.M.J. helped performed histological stains and data analysis. E.R. designed the AAV-TTR-FGF15 construct. (This needs to be adjusted based on final manuscript content).

## Supplementary Figures and Tables

**Supplementary Figure 1.**
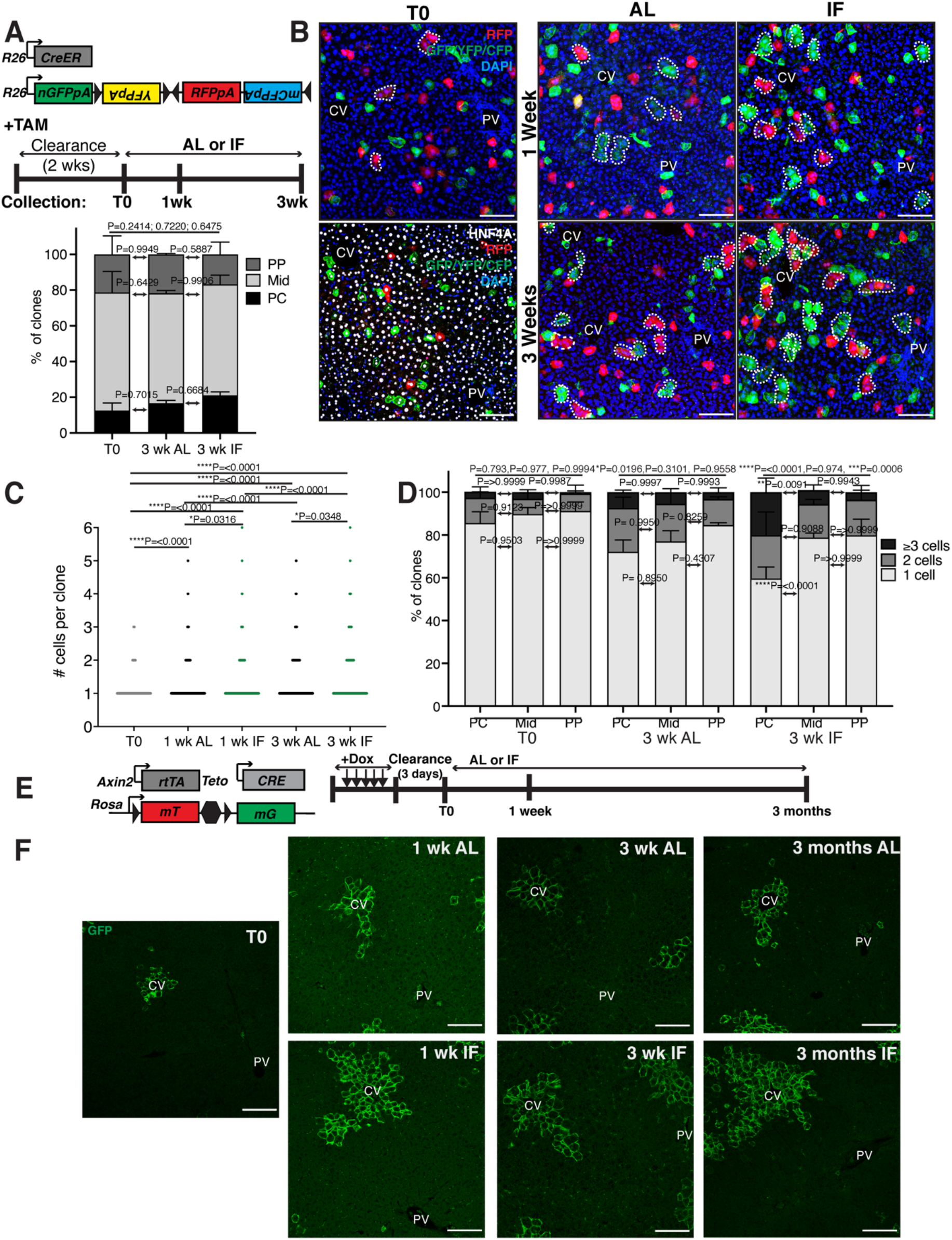
Hepatocyte proliferation kinetics in ad libitum fed and intermittent fasted animals. **(A)** Schematic of unbiased system to trace cell proliferation during ad libitum feeding (AL) and intermittent fasting (IF). *R26-CreER* mice were crossed to *R26-Confetti* mice enabling permanent cell labeling and lineage tracing by 4 fluorescent reporters after tamoxifen administration. **(B)** At TO, mostly single, HNF4A+ hepatocytes were labeled throughout the lobule. At 1 week and 3 weeks, multicellular hepatocytes clones (dotted circles) grew in AL and IF livers, with increased pericentral growth in IF. **(C)** Number of hepatocytes per 3D clone at each collection in A. Mann-Whitney test. 439-602 clones analyzed at TO, 430-1085 clones at 1 wk AL, 625-904 clones at 1 wk IF, 523-615 clones at 3 wk AL, and 442-615 at 3 wk IF. N=3. **(D)** Percentage of 3D clones consisting of different cell sizes, from C, in different liver lobule locations. PC, pericentral; Mid, midlobular; PP, periportal. Differences between 1-, 2- and >3 cell clones in periportal and pericentral zones are indicated by *P* values above bars. 2-way ANOVA. N=3.**(E)** Schematic of system to trace hepatocyte proliferation over 3 months of IF or AL treatment. *Axin2-rtTA*; *Teto-Cre*; *R26-mTmG mice* were induced with tamoxifen enabling permanent cell labeling and lineage tracing by reporter GFP.(F) GFP immunofluorescent images highlightling increase in hepatocyte proliferation from TO, 1-week, 3-weeks and 3-months after AL or IF treatment. \*\*\*\*\**P*<0.0001, \*\*\**P*<0.001, \*\**P*<0.01, \**P*<0.05. Error bars indicate standard deviation. Scale bar, 100μm.

**Supplementary Figure 2.**
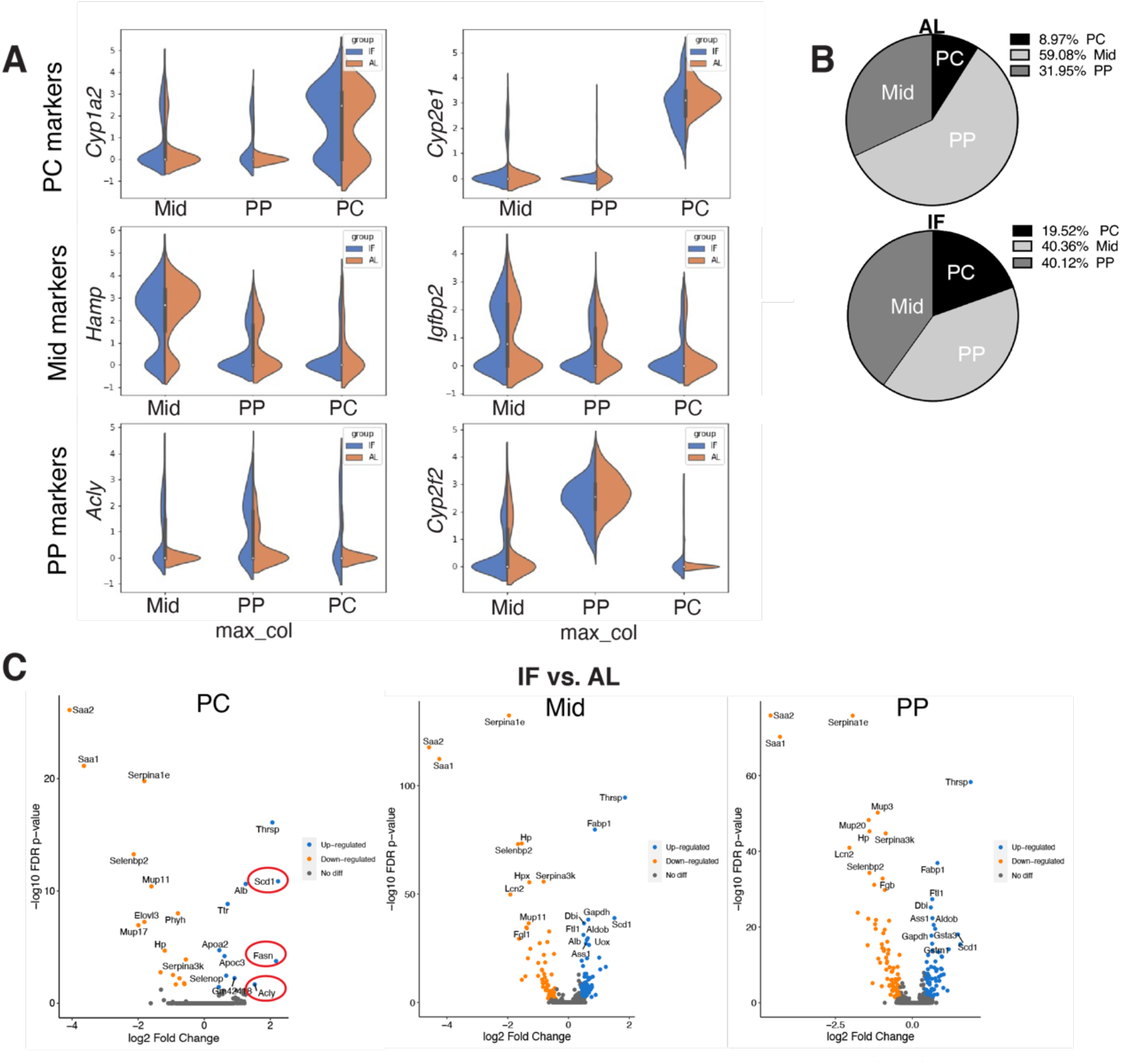
Single Cell RNA-seq comparing hepatocytes in ad libitum fed and intermittent fasted livers. **(A)** Violin plots, from scRNA-seq, demonstrating zonal marker gene expression used to classify single hepatocytes from AL and IF livers as PC, Mid and PP hepatocytes. **(B)** Pie charts of hepatocyte zonal populations identified in scRNA-seq highligting increase in PC hepatocytes in IF compared to AL livers. **(C)** Volcano plots highlighting differentially expressed transcripts in IF vs. AL livers in PC, Mid and PP hepatoyctes. Red circles highlighting transcripts invovled in de novo lipogenesis.

**Supplementary Figure 3.**
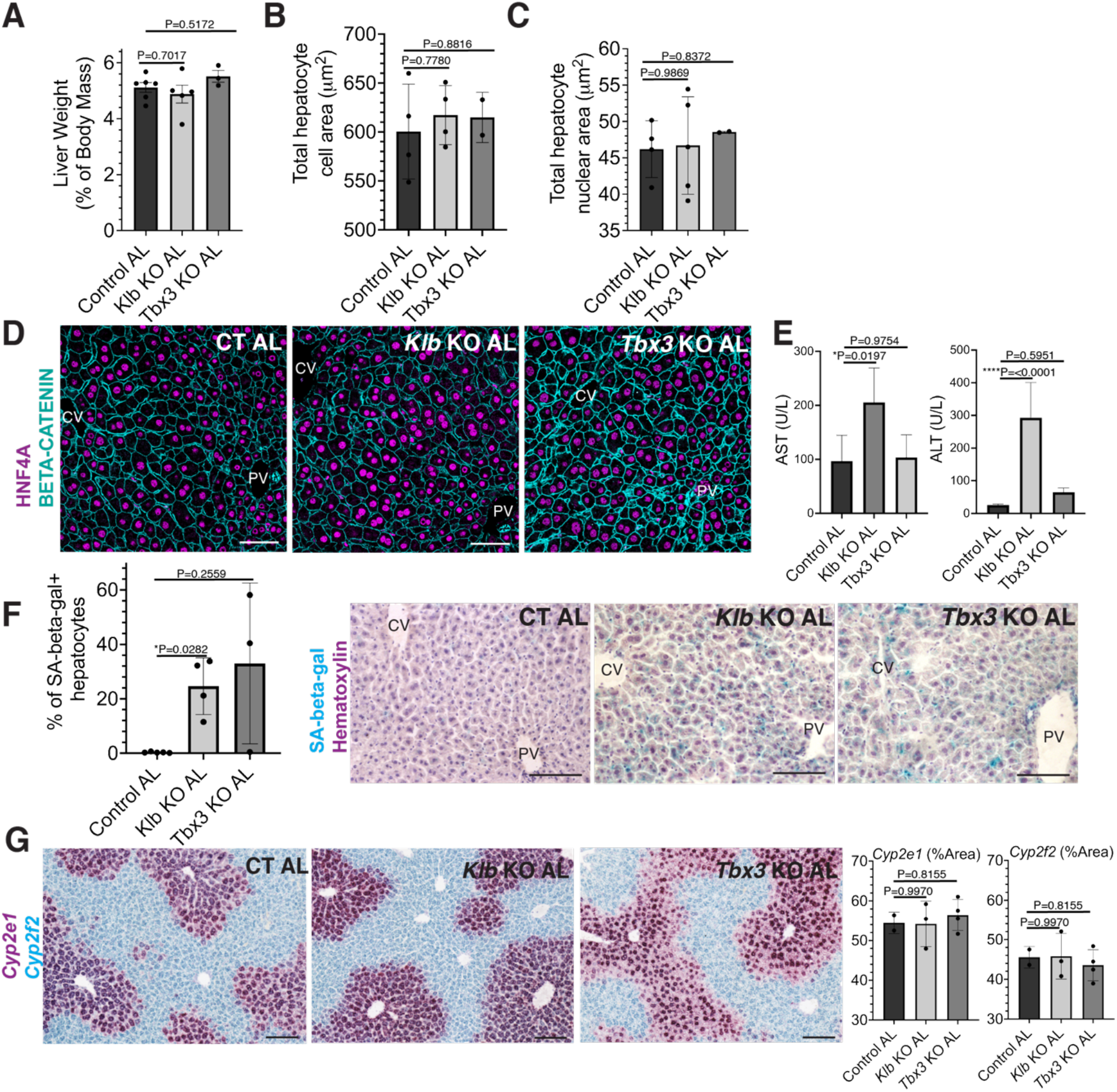
Short-term loss of *Tbx3* or *Klb* does not disrupt the hepatostat during ad libitum feeding. **A-H** AL *Klb* KO, AL *Tbx3* KO and CT AL livers were assessed at the same time point in Fig. 4 (3 weeks after ad libitum feeding). **(A)** Liver to body weight ratio. **(B-C**)Hepatocyte cell and nuclear area. **(D)** Immunofluorescence images for BETA-CATENIN and HNF4A highlighting hepatocyte cell and nuclear area during AL livers. **(E)** AST and ALT liver injury marker presence in serum. **(E)**Quantification and representative images ofsenescence-associated beta-galatosidase stains. **(G)** RNAscope for pericentral marker *Cyp2e1* and periportal marker *Cyp2f2*. All statistics were performed on 3-5 animals using 1-way ANOVA. \*\*\*\**P*<0.0001, \**P*<0.05. Error bars indicate standard deviation. Scale bar, 100μm.

**Supplementary Figure 4.**
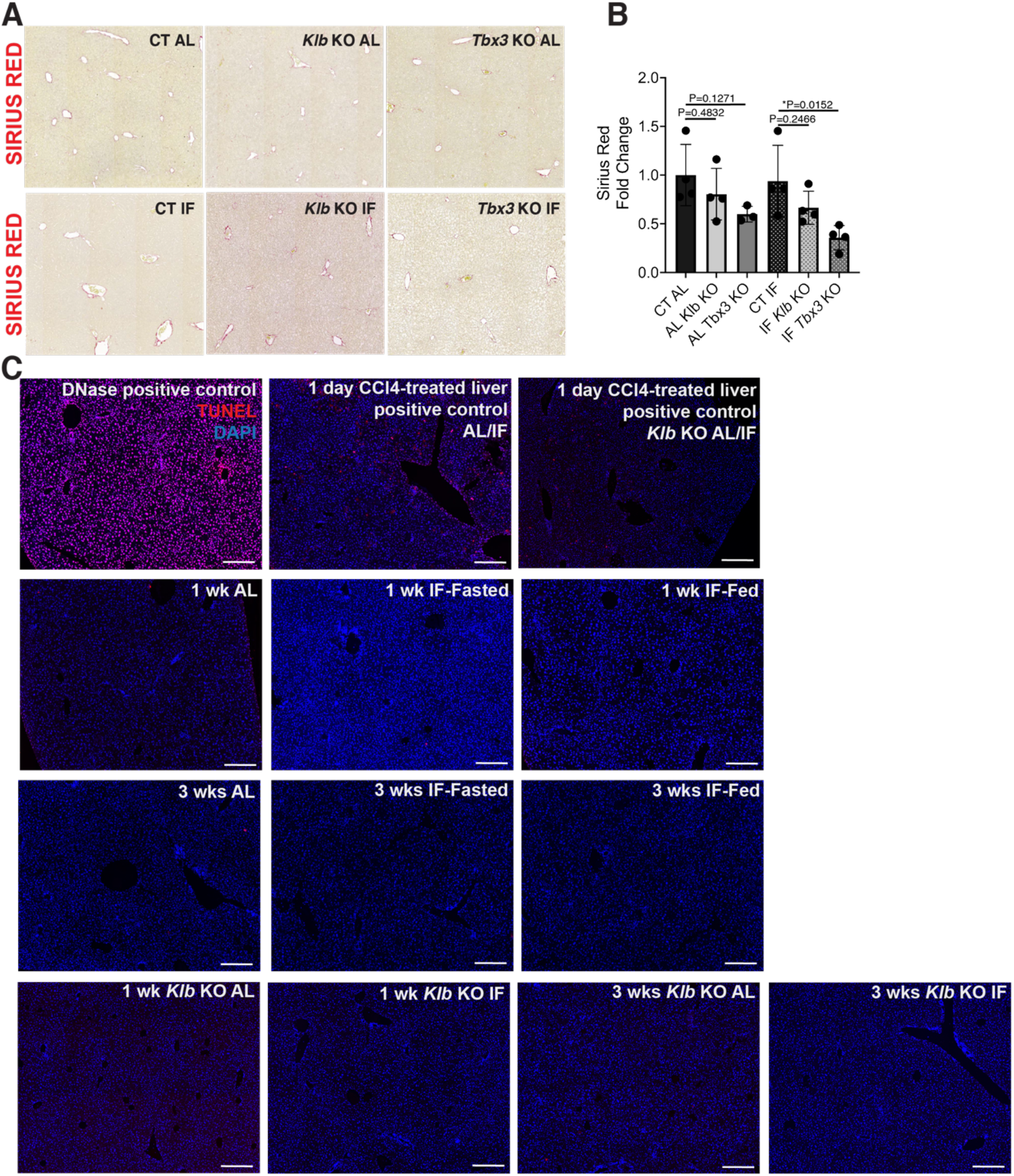
Fibrosis and cell death assessment of Control, *Klb KO* and *Tbx3* KO IF and AL-treated livers. **(A-B)** Sirius red staining and quantification to assess for liver fibrosis. **(C)** TUNEL stains on livers. All statistics were performed on 3-5 animals using 1-way ANOVA. \**P*<0.05. Error bars indicate standard deviation. Scale bar, 100μm.

**Table S1.**
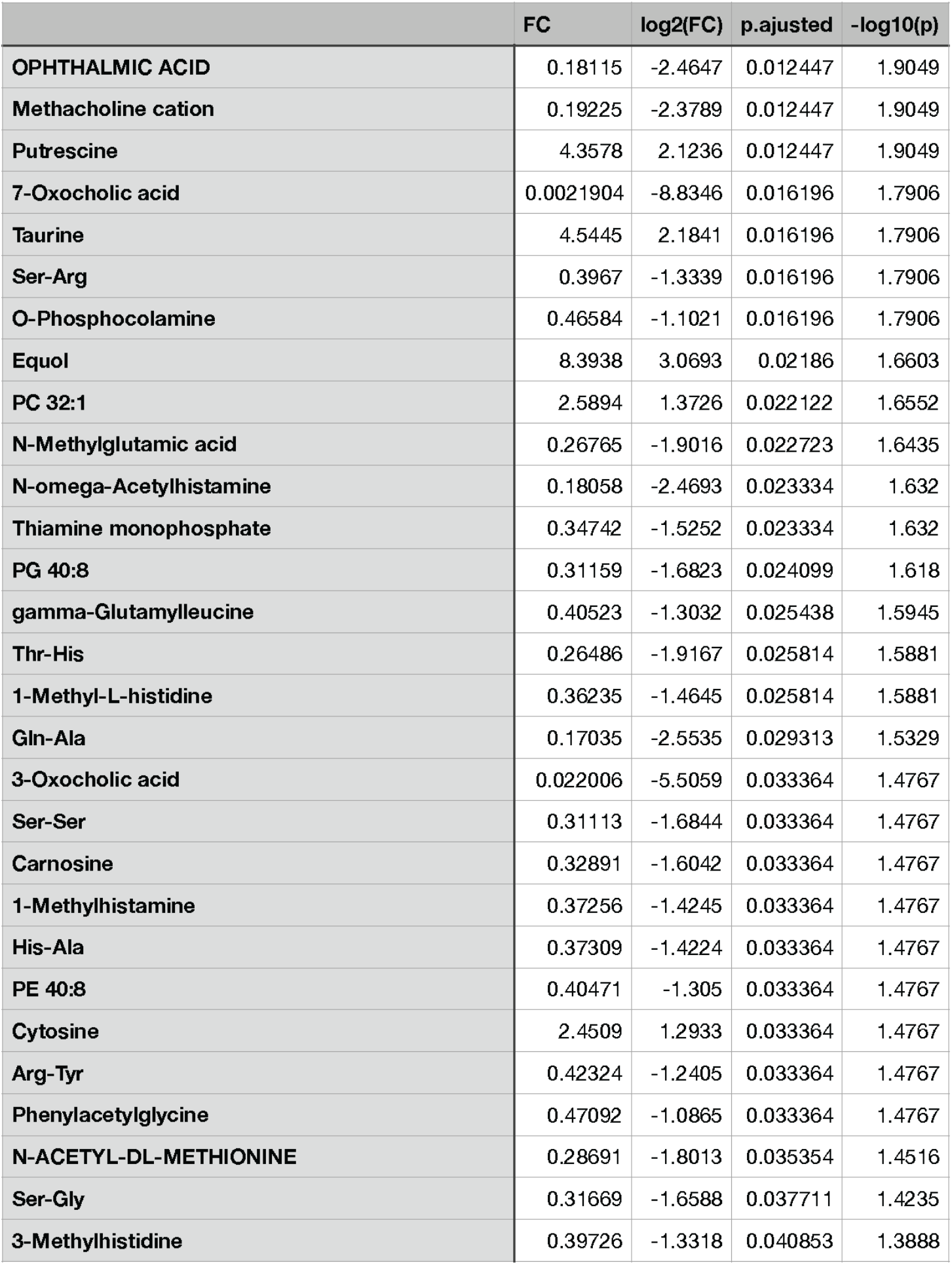
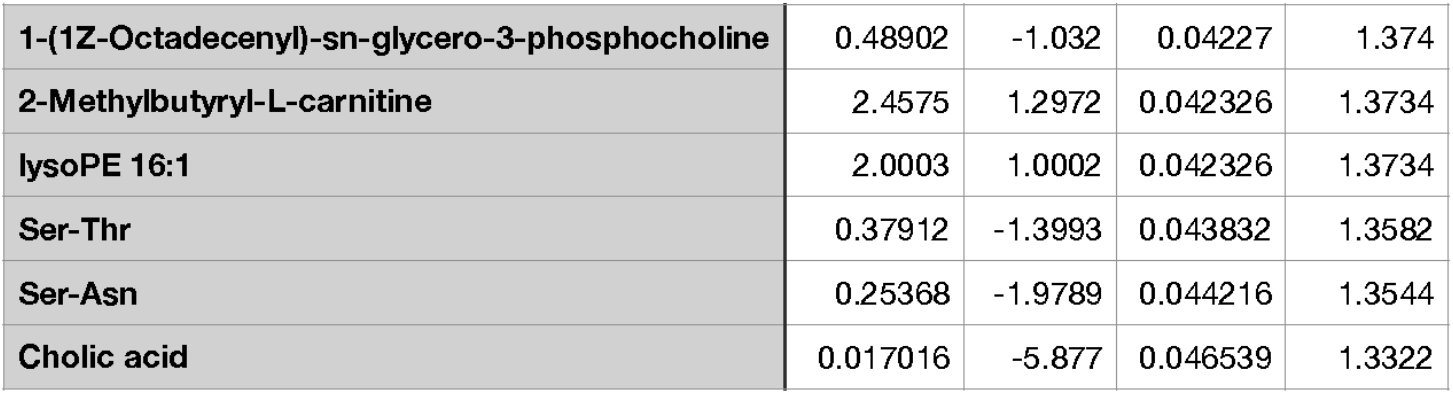
Metabolomics IF vs. AL Significantly Changed

